# Genome-wide DNA methylation and multi-omics study of human chondrocyte ontogeny and an epigenetic clock analysis of adult chondrocytes

**DOI:** 10.1101/2021.08.02.454544

**Authors:** Arijita Sarkar, Siyoung Lee, Ruzanna Shkhyan, Nancy Q. Liu, Ben Van Handel, Jenny Magallanes, Youngjoo Lee, Litao Tao, Neil Segil, Jason Ernst, Steve Horvath, Denis Evseenko

## Abstract

Articular chondrocytes undergo functional changes and their regenerative potential declines with age. Although the molecular mechanisms guiding articular cartilage aging is poorly understood, DNA methylation is known to play a mechanistic role in aging. However, our understanding of DNA methylation in chondrocyte development across human ontogeny is limited. To better understand DNA methylome changes, methylation profiling was performed in human chondrocytes. This study reveals association between methylation of specific CpG sites and chondrocyte age. We also determined the putative binding targets of STAT3, a key age-patterned transcription factor in fetal chondrocytes and genetic ablation of STAT3 induced a global genomic hypermethylation. Moreover, an epigenetic clock built for adult human chondrocytes revealed that exposure of aged adult human chondrocytes to STAT3 agonist, decreased epigenetic age. Taken together, this work will serve as a foundation to understand development and aging of chondrocytes with a new perspective for development of rejuvenation agents for synovial joints.

## Introduction

Tissue regeneration occurs widely in the animal kingdom ^1^. However, regenerative potential varies greatly across animals. Invertebrates and phylogenetically lower vertebrates, such as salamanders and zebrafish, often possess a higher regenerative capacity, and are capable of regenerating substantial parts of their body ^2^. In contrast, mammals have a very limited regenerative capacity. Articular chondrocytes have very limited potential for intrinsic healing and repair ^3^. Loss and degradation of articular chondrocytes is a significant cause of musculoskeletal morbidity. With aging, the regenerative potential of chondrocytes decreases with significant changes in mechanical, structural, matrix composition, and surface fibrillation ^4^. Although, the cellular and molecular mechanisms for chondrocyte regeneration are poorly understood, it is believed to be a cumulative combination of many molecular pathways.

Recent studies in this field have determined the importance of epigenetic regulation in mediating the process of aging ^5^. DNA methylation is a crucial player for epigenetic regulation of aging ^6, 7^. It is a biochemical process characterized by gain of methylation at the fifth carbon of cytosines i.e., 5-methylcytosine and occurs predominantly in cytosines followed by guanine residues (CpG). DNA methylation has diverse roles in several mammalian developmental stages, including genomic imprinting and X-chromosome inactivation ^8^ and is mediated by DNA methyltransferases. Although CpG methylation across mammals is tissue-specific, nearly 70-80% of CpGs in the mammalian genome are methylated. Establishment and regulation of DNA methylation is dynamic and varies considerably between different developmental stages and ages^9^. Although the mechanisms that drive changes in the methylome during aging are not well understood, but they have been attributed to environmental and spontaneous epigenetic changes^10^. Because DNA methylation changes are reversible, they are an attractive therapeutic target for aging. Previously, molecular markers like telomere length ^11^ and gene expression ^12^ were used to predict age across various tissues and organisms. However, with the advent of genome-wide methylation profiling, methylation pattern changes in CpG sites have been used to predict the biological age of individuals ^13^. The dynamics of methylation in aging have impelled researchers to develop ‘epigenetic clocks’ as the new standard to accurately predict biological age ^14, 15^. However, the impact of DNA methylation on chondrocyte development across human ontogeny has not been studied to date.

STAT3 is a well-known master transcriptional factor that exhibits a repertoire of signaling pathways in various tissues and contexts ^16, 17^, including self-renewal, proliferation, and pluripotency ^18, 19^. STAT3 also regulates chromatin accessibility via DNA methyltransferases ^20, 21^ and histone modifiers ^22^. Our recent studies have shown that STAT3 is highly activated in developing fetal chondrocytes ^23^. Moreover, the levels of active phosphorylated STAT3 (pSTAT3) are higher in fetal as compared to adult chondrocytes ^23^. However, the binding targets of STAT3 in human chondrocyte ontogeny and their potential role in maintaining the immature phenotype of fetal chondrocytes via epigenetic regulation has not been explored.

Thus, in this work, we study the dynamic genome-wide methylation profile of human chondrocytes across ontogeny. We have determined correlation between methylation of specific CpG sites and chondrocyte age. We also investigate the enrichment of chromatin states in these age-correlated CpGs. Besides, we also explored the putative binding targets of STAT3, a key age-patterned TF in fetal chondrocytes along with impact of STAT3’s genetic manipulation on genome-wide DNA methylation. Moreover, we apply a novel epigenetic clock for adult human chondrocytes that accurately predicts epigenetic age. We utilized this clock to gain further insight into the effect of a small molecule STAT3 agonist in decreasing epigenetic age of aged adult chondrocytes. In a nutshell, these findings will serve as a foundation to understand the global DNA methylation profile of human chondrocytes and help develop new therapeutic interventions to reverse or slow down aging.

## Results

### Epigenome-wide association study (EWAS) identifies age-correlated CpGs in non-cultured human fetal and adult chondrocytes

We performed DNA methylation profiling for non-cultured human fetal (n=8) and adult chondrocytes (n=22) and identified regulatory genes associated with ontogeny specification. The DNA methylation ß-values across all samples (**Fig 1a**) from 865,859 CpG sites follows a bimodal distribution with peaks around 0 (unmethylated) and 1 (methylated). Evaluation of global methylation patterns (hypomethylation and hypermethylation) across the ontogeny revealed correlation with chondrocyte age (**Table S1**). Further site-specific genome-wide pattern of DNA methylation (**Fig 1b-c**) showed a predominant proportion of age-correlated CpG sites to be statistically significant (p-value<0.05). These CpGs showing either gain or loss of methylation across ages (i.e., hypermethylated or hypomethylated respectively) were not evenly distributed across the genome, showing prevalence in open sea regions and mostly confined in the gene body (**Fig 1d**). Several chondrocyte-associated genes including UCMA, SOX11, BMPR1B, CSPG4, COL2A1, ITGA10, COL9A1, and RUNX2 are known to be expressed during development ^24^. We thus explored the methylation level for all age-correlated CpG probes associated with these chondrogenic genes (**Fig S1**). Our data suggests that with aging, age-correlated CpGs associated with chondrogenic genes gain methylation (**Fig 1e**) and show expression downregulation as revealed by the transcriptomic ^24^ and single cell sequencing data ^25^ for non-cultured fetal and adult chondrocytes (**Fig 1f-g**). Besides, age-correlated CpGs losing methylation with age and the transcriptional profile for the associated genes has been shown in **FigS2-3.** We also examined the methylation status of all age-correlated CpG probes associated with microRNA (miRNA) genes (**Fig S4**), which are known to play an important role in chondrocytes during development ^26–29^. The age-correlated CpGs for these miRNAs also gain methylation with age and show downregulation in adult chondrocytes as revealed by the miRNA-sequencing data (**Fig 1h-i, Table S2**). miRNAs associated with age-correlated CpGs losing methylation with age has been shown in **FigS5.** Overall, age-correlated CpGs, show a distinct methylation profile in fetal and adult chondrocytes, which in turn governs the ontogeny-specific phenomenon of development.

**Figure 1.**
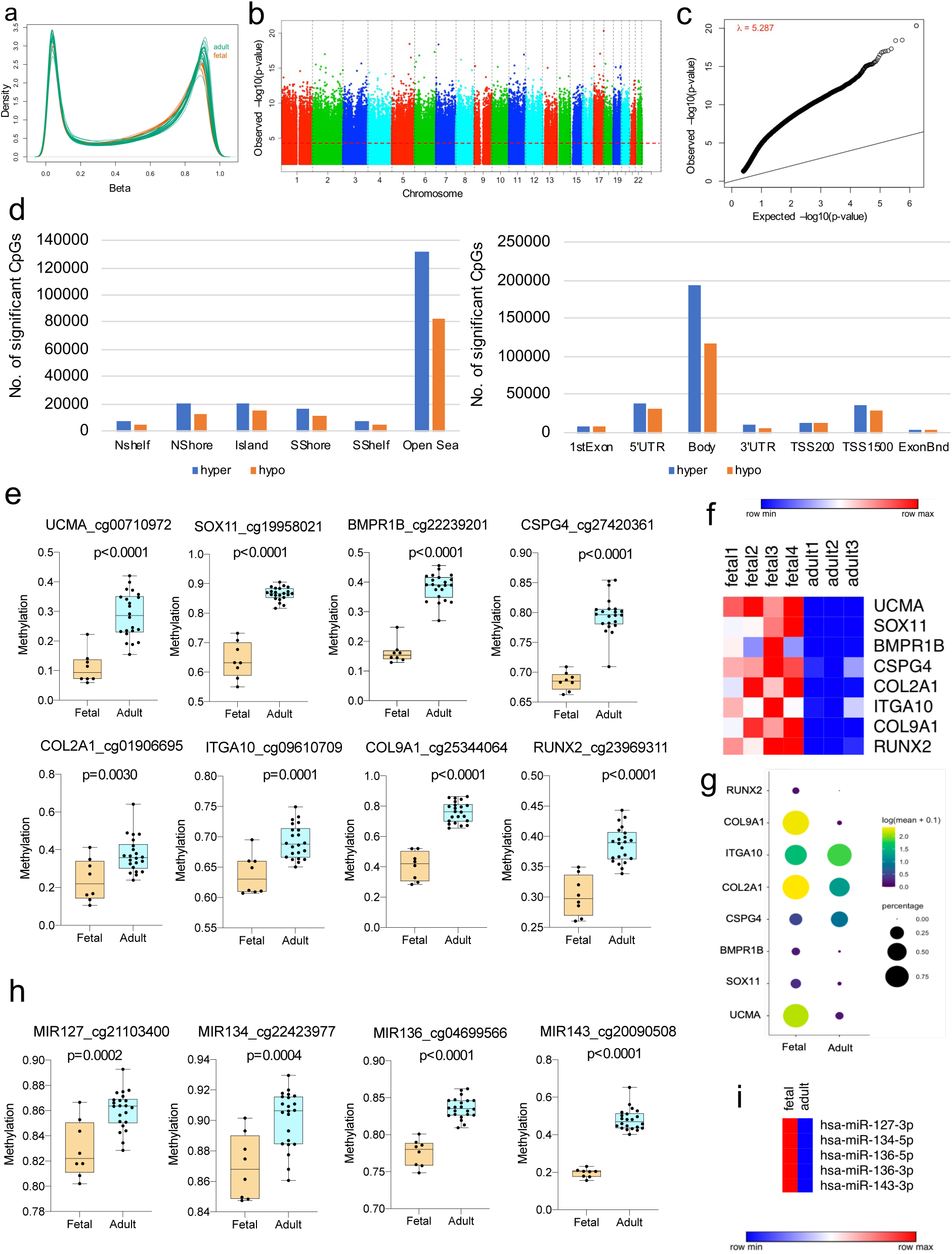
Epigenome-wide association study for non-cultured fetal and adult chondrocytes. **a.** Density plot for all samples. CpGs are shown for 865,859 loci. **b**, Manhattan plot showing chromosomal locations of age-correlated CpGs along with −log_10_(*P* values) for association at each locus. The red dotted line indicates the p-value threshold of 0.05. **c.** QQ plot showing observed versus expected −log_10_(*P* values) for age-correlated CpGs **d.** Distribution of CpG features among age-correlated CpGs. hyper= CpGs gaining methylation with age, hypo= CpGs losing methylation with age **e.** Boxplot showing methylation level of representative age-correlated CpGs (i.e., CpGs with highest hypermethylation change across age) corresponding to chondrogenic genes. **f.** Transcriptomic profile for the chondrogenic genes (shown in e) in fetal and adult chondrocytes. **g.** Dot plot showing expression for the chondrogenic genes (shown in e) from single cell sequencing in fetal and adult chondrocytes **h.** Boxplot showing methylation level of representative age-correlated CpGs (i.e., CpGs with highest hypermethylation change across age) corresponding to miRNAs expressed in fetal and adult chondrocytes. **i.** miRNA expression profile for the miRNAs shown in h. Hinges of all boxplots extend from the 25th to 75th percentiles. The line in the middle of the box is plotted at the median. P-values are calculated using 2-tailed Student’s t test.

### Age-correlated CpGs are associated with distinct chromatin signatures

It has been previously reported that DNA methylation patterning is governed by various chromatin states such as histone modifications, and nucleosome positioning ^30^. Additionally, various chromatin remodeling factors might interact with DNA methyltransferases, guide them to specific DNA sequences and modulate transcriptional activation/repression. A closer inspection into the genes associated with the age-correlated CpGs revealed enrichment of Gene Ontology terms involving binding and activity of several histone modifiers including enhancer-mediated binding (**Fig 2a**). Thus, we hypothesized that age-correlated CpGs might be associated with distinct chromatin states in chondrocytes. Accordingly, we determined the chromatin states associated with age-correlated CpGs (i.e. both hypermethylated(204549 CpGs) and hypomethylated(132383 CpGs)) in fetal and adult chondrocytes using the ChromHMM chromatin state model previously generated by our group ^24^ based on data from four histone modifications (H3K4me3, H3K27me3, H3K4me1, and H3K27ac)(**Fig 2b**). We observed that CpGs in fetal chondrocytes, which gain methylation with age, show stronger enrichment for a poised promoter or bivalent state, characterized by the co-existence of both activating (H3K4me3) and repressing (H3K27me3) marks. Interestingly, bivalent chromatin states has been previously known to be enriched in developmentally important genes ^31^. Besides CpGs in adult chondrocytes, which lose methylation with age are most enriched for the active enhancer chromatin state suggestive of transcriptional regulation from these regions. Of note, gain or loss of methylation in CpGs correlated with age in both fetal and adult chondrocytes show enrichment for chromatin states associated with enhancers (marked by H3K27ac) which might indicate the previously known fact that chondrocytes acquire cell-type-specific enhancers upon differentiation ^32^. We further investigated the chromatin state for the chondrogenic genes mentioned previously in Fig 1e and closer inspection of these loci demonstrate presence of active histone modifications characterized by presence of H3K27ac while H3K27me3 repressive mark is mostly absent (**Fig 2c**). Taken together, these findings affirm that age-correlated CpGs are intrinsically tied to chromatin state and corroborate with regulation of chondrogenic genes as shown previously using methylation and transcriptomic data for fetal and adult chondrocytes.

**Figure 2.**
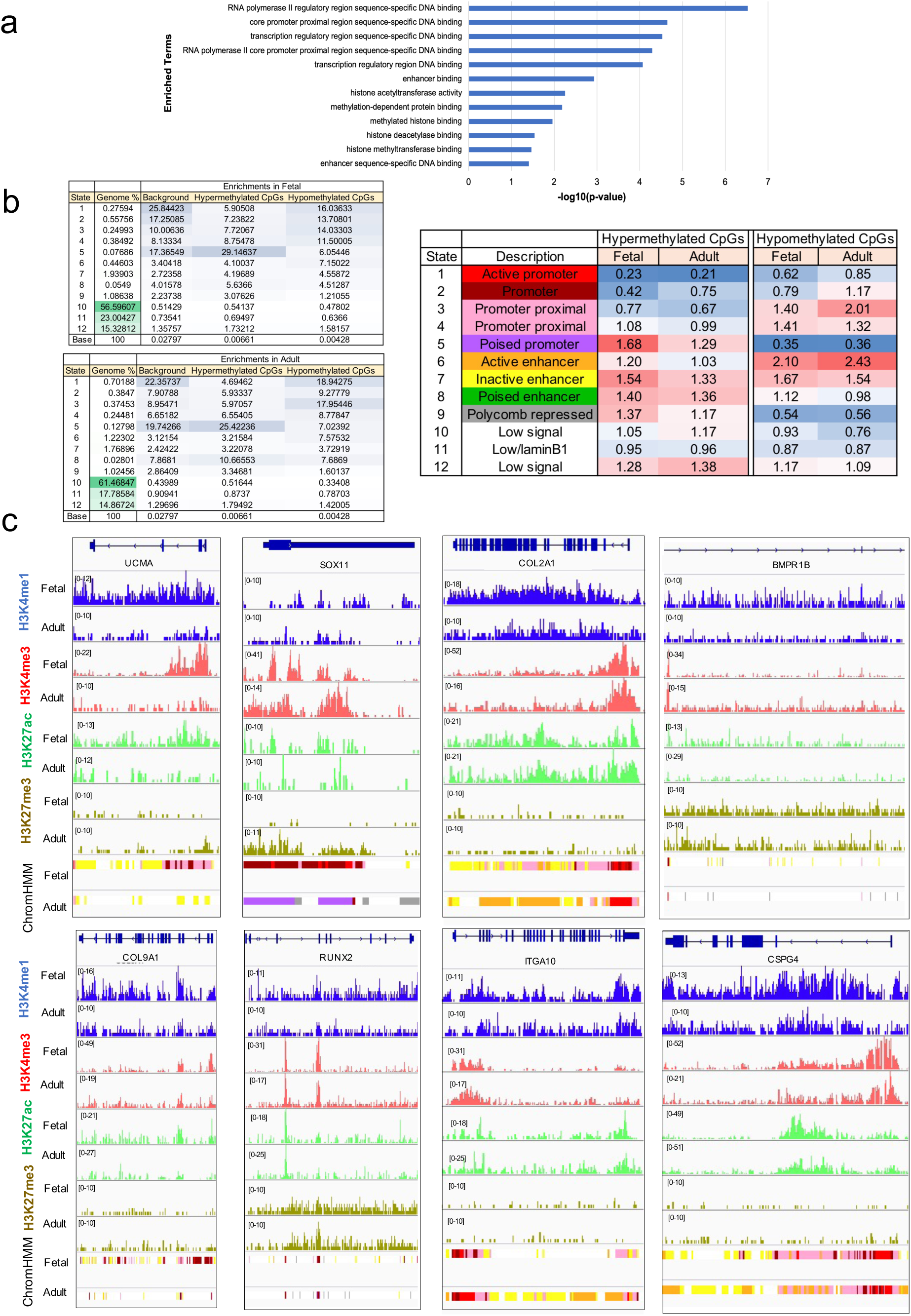
Age-correlated CpGs are associated with distinct chromatin states. **a.** Gene ontology analysis for genes associated with the age-correlated CpGs show enrichment for terms associated with chromatin states and histone modifications using Fisher’s exact test (p-value<0.05) **b.** ChromHMM model shows enrichment of the 12 chromatin states for age-correlated CpGs in fetal and adult chondrocytes. Hypomethylated CpGs refer to CpG sites which are losing methylation with chondrocyte age. Hypermethylated CpGs refer to CpG sites which are gaining methylation with chondrocyte age. Emission probabilities (left panel) shows the occurrence of CpGs in each chromatin state. Rows correspond to chromatin states. The occurrence of CpGs in each chromatin state is represented by color code: 0(white) to 100(blue). Chromatin state enrichments (right panel) shows the enrichment score for CpGs in each chromatin state. A 3-color code was used to represent the range of enrichment score: Lowest value(blue), 50percentile(white) and Highest value(red). **c.** Chromatin data for chondrogenic genes shown in fetal and adult chondrocytes. ChromHMM tracks are colored according to the chromatin state color code in b (right panel).

### Genome-wide putative STAT3 targets differ in development and disease

STAT3 exhibits a plethora of functions with context-specific roles in skeletal development, inflammation, and neoplastic growth ^33^. It is also involved in regulating methylation of CpGs sites by interacting with DNA methyltransferases ^20^. Also as mentioned previously our lab has observed STAT3 to be highly expressed in fetal chondrocytes in comparison to healthy adults ^23^. Besides, pSTAT3 is also highly expressed in osteoarthritic chondrocytes in comparison to healthy adults (**Fig S6**). Hence, it is quite evident that although STAT3 is highly expressed in fetal and osteoarthritic chondrocytes when compared to healthy adults, the outcomes downstream of STAT3 are different in each context. This led us to hypothesize that STAT3 has different context-specific transcriptional targets that differ in development and disease.

To gain further insight into the context-specific putative targets of STAT3, we performed Cleavage Under Targets and Release Using Nuclease (CUT&RUN) ^34^ profiling for fetal, adult, and osteoarthritic chondrocytes (n=2 for each case). The average profile plot for peaks shows binding around the transcription start site (TSS) and extending to genic regions with confidence intervals shown by the shadows following each curve. Confidence intervals were estimated by bootstrap method using 500 iterations (**Fig 3a**). Heatmaps centered around the peak summits shows enrichment of reads (**Fig 3b**). Most of the STAT3-binding sites were located in the distal intergenic regions, suggesting STAT3 might regulate the expression of its putative targets by binding to distal regulatory elements (**Fig 3c)**. Interestingly, epigenetic regulation mediated by STAT3 via binding to intergenic regions has been reported previously ^35, 36^. Further, gene enrichment analysis for putative STAT3 binding targets revealed distinct pathways and molecular functions regulated in fetal and adult chondrocytes (**Fig 3d)**. For instance, the Wnt signaling pathway, which is enriched in fetal chondrocytes, is known to maintain an immature phenotype by regulating self-renewal and pluripotency in human pluripotent stem cells ^24, 37^. In contrast, enrichment of extracellular matrix (ECM) receptor interaction in adult chondrocytes is suggestive of the gradual degradation of ECM with age ^38^. We next identified the enriched DNA motifs present in the putative STAT3 targets for both fetal and adult chondrocytes (**Fig 3e**). For fetal chondrocytes, we obtained motifs from several well-known and important transcription factors known to modulate early development, including SOXs (SOX4, SOX6) ^39^ and LEF1^40^. Similar analysis for adult chondrocytes showed enrichment for GATA1, GATA2, IRF4, GLI3, CTCF binding motifs. Although the role of these genes in chondrocytes remains unclear, these transcription factors are known to be essential for differentiation and lineage commitment in different cell types ^41–45^. To date, researchers have uncovered several STAT3 binding targets across various other tissues and cell types. Since STAT3 binding targets have not been studied in human chondrocytes, we were interested in exploring the exclusive putative binding targets in human chondrocytes. Thus we overlapped the STAT3 targets reported till date in ChIP-Atlas ^46^ and CistromeDB ^47, 48^ with our analysis (**Fig 3f**). Interestingly, we obtained 1858 exclusive targets in human chondrocytes (**Table S3**).

**Figure 3.**
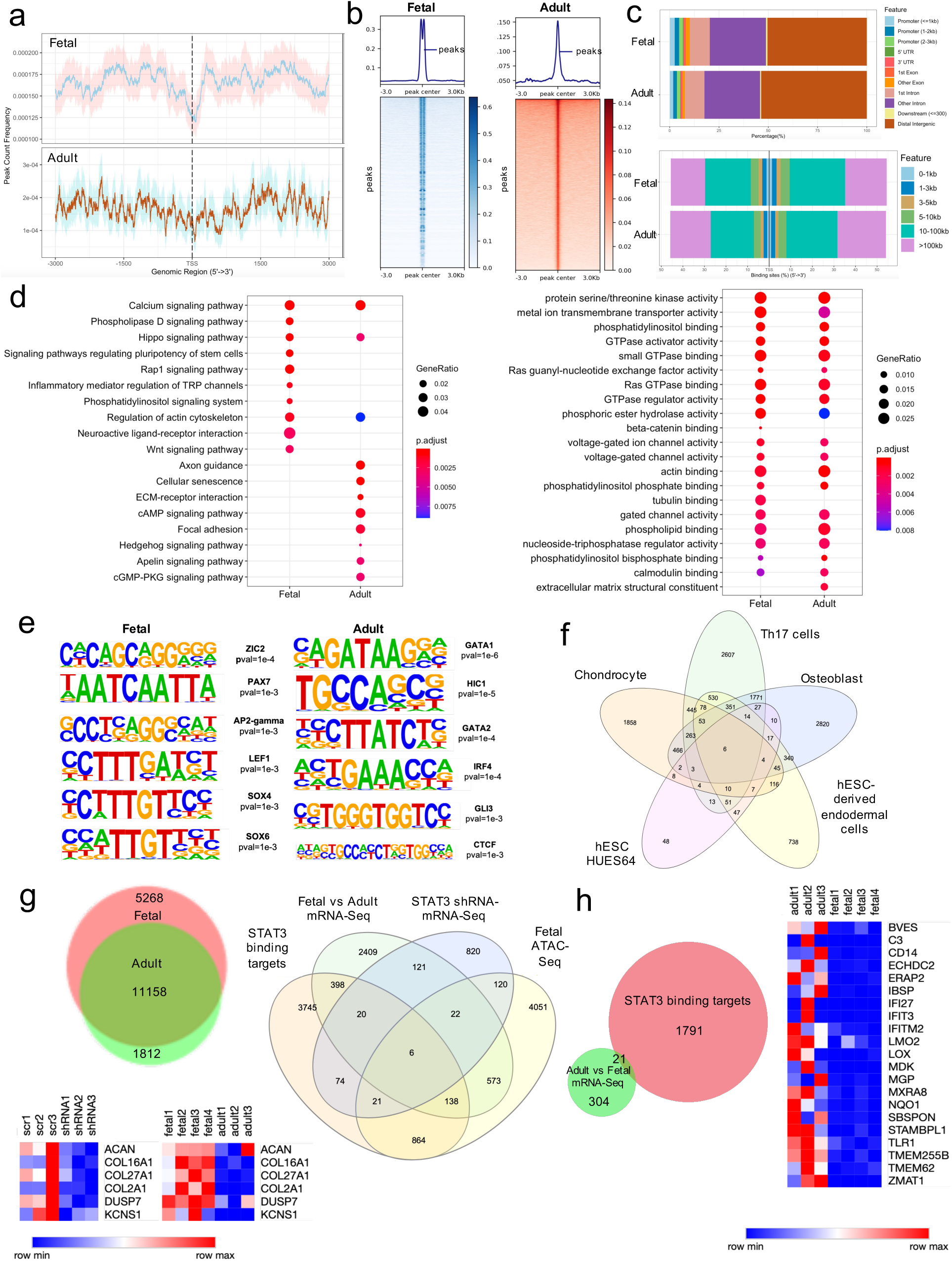
STAT3 binding targets during chondrocyte development. **a.** Distribution of peak count frequency across ±3kb of TSS. Confidence intervals shown by the shadows following each curve were estimated by bootstrap method using 500 iterations **b.** Heatmap showing enrichment of reads in peak summits. **c.** Bar plot showing the distribution of genomic features for peaks in fetal and adult chondrocytes. **d.** Gene enrichment analysis of putative STAT3 target genes. P-values were adjusted using Benjamini-Hochberg correction method **e.** DNA motif enrichment analysis for putative STAT3 binding targets. Binomial distribution was used to score motifs. **f.** Chondrocyte specific putative STAT3 binding targets compared to other tissue types. **g.** Venn diagram showing the overlap between putative STAT3 targets in fetal and adult chondrocytes. 5268 exclusive fetal chondrocyte targets were overlapped with Fetal vs adult mRNA-seq, STAT3 shRNA mRNA-seq and Fetal ATAC-seq data. Heatmaps show the expression profile (STAT3 knocked-down fetal chondrocytes and fetal vs adult chondrocytes) of the 6 final targets obtained for fetal chondrocytes **h.** 1812 exclusive adult chondrocyte targets were overlapped with Fetal vs adult mRNA-seq. Heatmaps show the expression profile (fetal vs adult chondrocytes) of the 21 final targets obtained for adult chondrocytes.

We overlapped the putative binding targets obtained for fetal and adult chondrocytes and determined targets exclusively present in fetal chondrocytes. To evaluate the concordance between these fetal chondrocyte exclusive 5268 putative STAT3 targets and gene expression (**Fig 3g**), we compared them to transcriptomics data from i) fetal and adult chondrocytes ^24^ and ii) STAT3 knocked down fetal chondrocytes ^49^. We also performed ATAC-seq on fetal chondrocytes(n=3) (**Fig S7**) to check for chromatin accessibility. We obtained 6 well-known genes (ACAN, COL16A1, COL27A1, COL2A1, DUSP7, KCNS1) which had putative open chromatin regions. Interestingly, COL2A1 which is a key structural gene and plays a critical role in matrix anabolism was shown to have gained methylation with age (**Fig 1e, Fig S1)**. Upon a similar analysis with 1812 exclusive putative STAT3 targets in adult chondrocytes, we finally obtained 21 of them to be overlapping with transcriptomics data from adult chondrocytes (**Fig 3h**). Of these, CD14 and TLR1 have been shown to be losing methylation with age (**Fig S2-3)**.

Next, we assessed the role of STAT3 in disease by determining the putative binding partners in osteoarthritic chondrocytes by CUT&RUN and comparing them to those in development. As mentioned previously, STAT3 might regulate chondrocyte development and disease by binding to different partners dependent on context. The profile for osteoarthritic chondrocytes (**Fig S8a-c**) shows binding mostly in the distal intergenic region. We do observe that different pathways are regulated by STAT3 in the context of disease and development (**Fig S8d**). On motif analysis for the putative STAT3 binding sites we obtained DNA motif for NF-kB, which is a well-known transcription factor that mediates inflammation (**Fig S8e**). Recently, Wang et al. have demonstrated that STAT3 can speed up osteoarthritis through the NFkB signaling pathway ^50^. Other transcription factors that might regulate osteoarthritis via co-binding to STAT3 include TGIF1, JUNB, FOSL2 and FOXO1, mostly known for their role in inflammation ^51–54^. We next overlapped the putative targets obtained from fetal chondrocytes and osteoarthritic chondrocytes and determined the exclusive targets in disease. Of these 84 exclusive binding partners in disease, 16 targets were highly expressed in osteoarthritis in comparison to fetal chondrocytes as suggested by single cell sequencing data (**Fig S8f**). Thus, combinatorial analysis of this data provides critical insight into the multipotential, and context-specific mode of regulation exhibited by STAT3 during development and disease.

### Genetic manipulation of STAT3 induces global hypermethylation in fetal chondrocytes

Our lab has previously shown that STAT3 is essential for normal cartilage development and is highly expressed in anabolic fetal chondrocytes compared to healthy adult chondrocytes ^23^. Recently we have also shown that postnatal STAT3 deletion in 3-months-old mice lead to degradation of the growth plate ^49^. Moreover, upon STAT3 inhibition, an increase in apoptosis and decrease in proliferation was observed ^23^. In summary, STAT3 plays a predominant role in chondrogenesis, and its deletion leads to profound changes in early development. Thus, we hypothesized that genetic manipulation of STAT3 in fetal chondrocytes might have an impact on genome-wide DNA methylation. We transduced fetal chondrocytes with STAT3 shRNA (n=4) and scrambled (n=4) (**Fig S9**) and performed DNA methylation profiling. To understand the effect of STAT3 inhibition, we determined the differentially methylated CpGs. Density and volcano plots for the CpG sites suggested that 55697 CpGs are statistically significant (p-value<0.05) (**Fig 4a-b**). Interestingly, we found a significant number of CpGs have gained methylation (hypermethylated) in STAT3 knocked down fetal chondrocytes (**Fig 4c**). We strengthened our observation by looking into differentially methylated CpGs, that are correlated with age (**Fig 4d**). These CpG sites were unevenly distributed across the genome, and they were prevalent in the open sea region (**Fig 4d**). Differentially methylated CpGs which are age-correlated as well showed a significant increase in hypermethylation across the genome (**Fig 4e)**. Furthermore, we explored the concordance between genes associated with differentially methylated CpGs that gain methylation with age and transcriptomic data from i) fetal chondrocytes ^24^ and ii) STAT3 knocked down fetal chondrocytes ^49^ as well as STAT3 binding targets determined previously (**Fig 4f**). In summary, it can be concluded that upon STAT3 inhibition in fetal chondrocytes there is a global gain in methylation that might attribute to epigenetic aging of these cells.

**Figure 4.**
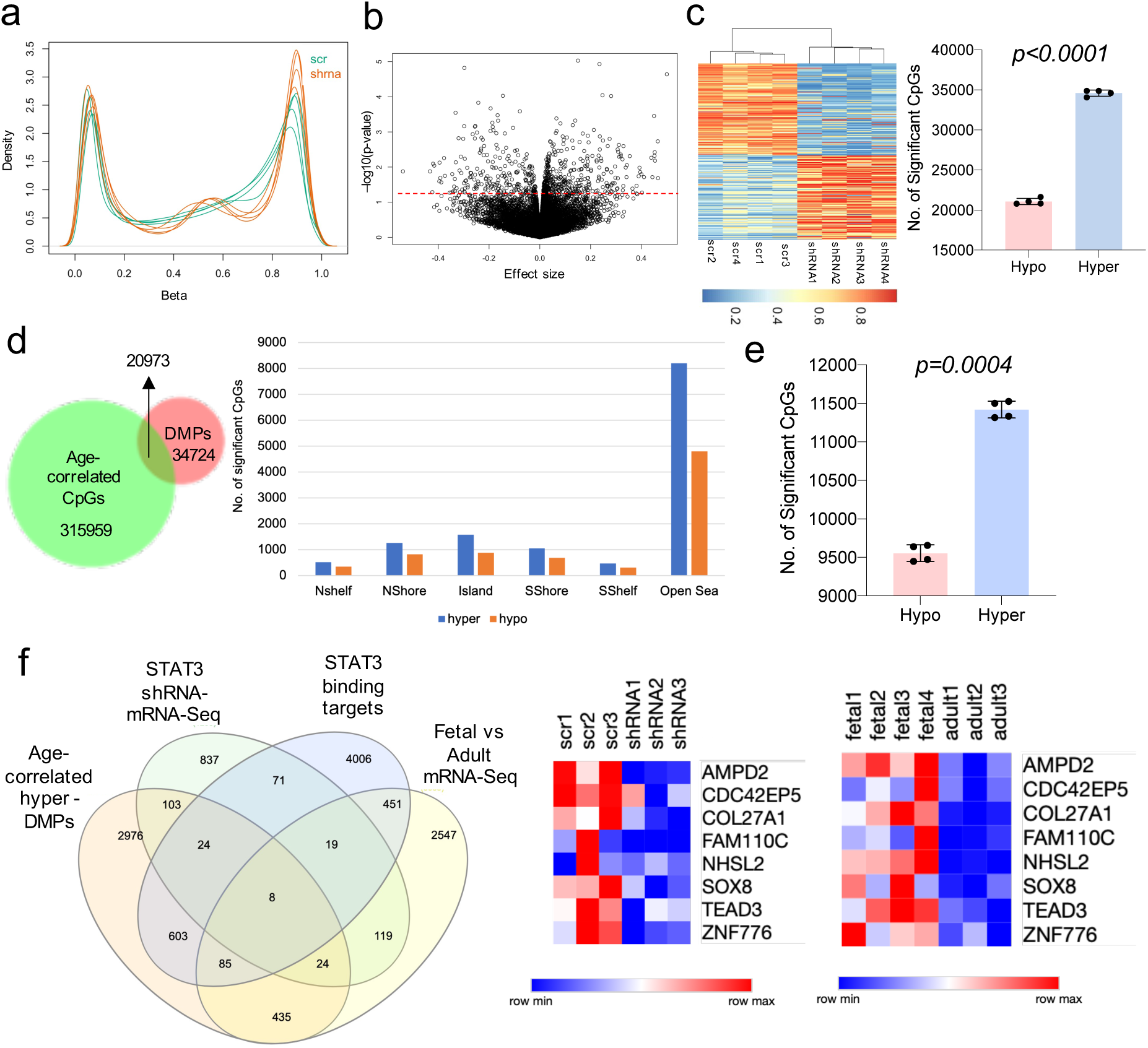
STAT3 knockdown induces genomic hypermethylation in fetal chondrocytes. DMPs= differentially methylated CpG probes. **a.** Density plot for all samples **b.** Volcano plot for all DMPs. Dotted red line indicates p-value threshold of 0.05. **c.** Heatmap showing the sample clustering based on DMPs. Bar diagram shows the gain in hypermethylation in DMPs. **d.** Venn diagram showing the overlap between DMPs and age-correlated CpGs. 20973 DMPs are age-correlated. Distribution of CpG features among these 20973 age-correlated DMPs. **e.** Bar plot shows gain in hypermethylation in age-correlated DMPs. **f.** Genes associated with age-correlated hypermethylated DMPs were overlapped with STAT3 shRNA mRNA-seq in fetal chondrocytes, putative binding targets in fetal chondrocytes and fetal vs adult mRNA-seq. Heatmaps show the expression profile (STAT3 knocked-down fetal chondrocytes and fetal vs adult chondrocytes) of the 8 genes. P-values are calculated using 2-tailed Student’s t test. Mean with standard deviation is plotted.

### A novel epigenetic clock for adult chondrocytes helps to accurately predict STAT3 agonist-induced global hypomethylation

Since the late 1960s, a vast majority of literature describes DNA methylation levels as having strong effects on the aging of tissues and cells ^55, 56^. DNA methylation based epigenetic clocks are the best biological age predictors till date ^57^. Several epigenetic clocks have been developed for various tissues across several species ^13^. To the best of our knowledge, we for the first time, have developed a novel epigenetic clock that is specific to human adult chondrocytes (**Fig 5a**). This clock utilizes DNA methylation data to estimate biological age of human adult chondrocytes with high accuracy (*r=0.97, p-value=2.4E-14*). Further, we used this novel clock to accurately predict epigenetic age of adult chondrocytes upon treatment with a STAT3 agonist.

**Figure 5.**
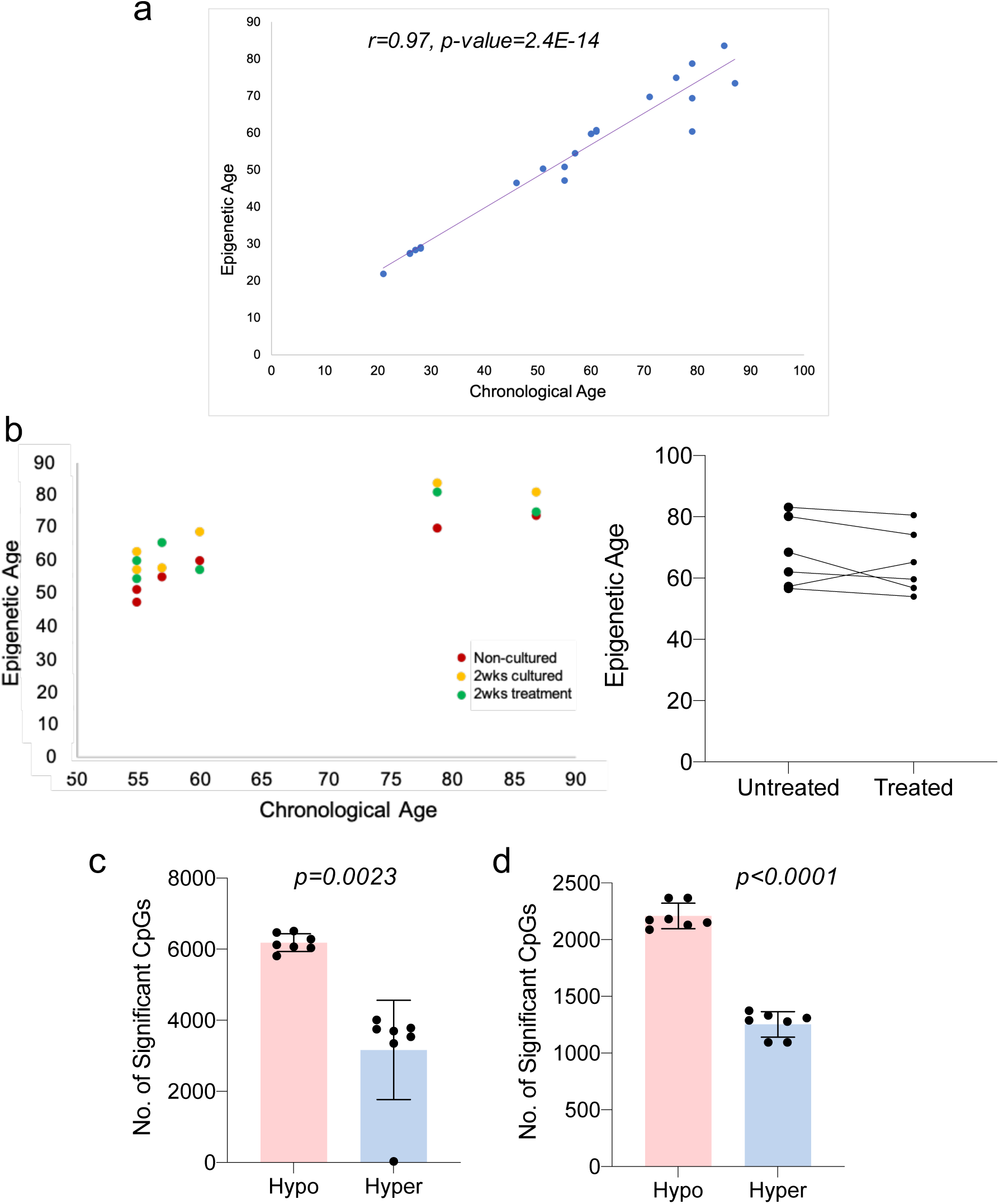
A novel epigenetic clock for adult chondrocytes. **a.** Epigenetic clock for adult chondrocytes shows high correlation between epigenetic age and chronological age. **b.** Administration of a small molecule STAT3 agonist to adult chondrocytes for 2 weeks lowers epigenetic age. **c.** Differentially methylated CpGs between 2wks cultured treated and untreated samples show global gain in hypomethylation **d.** Age-correlated differentially methylated CpGs between 2wks cultured treated and untreated samples show global gain in hypomethylation. P-values are calculated using 2-tailed Student’s t test. Mean with standard deviation is plotted.

Our lab previously performed a high throughput screening of 170,000 compounds and identified a small molecule which acts as a STAT3 agonist in adult chondrocytes, thereby reducing cartilage degeneration and structural damage ^23^. This small molecule increased proliferation while reducing apoptosis and hypertrophic responses in adult chondrocytes *in vitro*. Besides, this molecule was shown to promote cartilage repair in a rat osteochondral defect model with spontaneous healing in 4 weeks ^23^. Moreover, we have also shown that this compound plays a role in hair follicle stem cell activation via STAT3 ^58^. Hence, to gain further insight into the mechanism, we treated adult chondrocytes with or without STAT3 agonist for 2 weeks (n=6) and performed DNA methylation profiling. We hypothesized that treatment of adult chondrocytes with STAT3 agonist would make adult chondrocytes epigenetically younger. Interestingly, based on the novel clock, adult chondrocytes from 5 out of 6 tested donors showed a clear decrease in epigenetic age upon treatment for 2 weeks (**Fig 5b**). Thus, to strengthen our results, we determined the differentially methylated CpGs between 2 weeks cultured, treated and untreated samples and observed a global hypomethylation in treated samples (**Fig 5c**). We also evaluated the differentially methylated CpGs, which are age-correlated, and obtained global hypomethylation in treated samples (**Fig 5d**). Taken together these results suggest that pharmacological activation of STAT3 signaling in aged adult chondrocytes reduces their epigenetic age. These proof-of concept studies open a new perspective for development of rejuvenation agents for synovial joints.

## Discussion

Articular chondrocyte development and differentiation is governed by cell-specific gene expression patterns, which is in turn established and reinforced by DNA methylation ^59^. Here we generated a DNA methylation profile for human chondrocytes across ontogeny and determined the epigenome-wide changes in the methylome of fetal and adult chondrocytes. We showed association between methylation of CpG sites and chondrocyte age. Moreover, these age-associated CpGs are mainly confined to the open sea and gene body regions showing the distinct pattern of epigenetic regulation in chondrocytes. A closer inspection into the methylation pattern revealed gain of methylation with age in CpGs associated with chondrogenic genes. These observations were in concordance with upregulation of chondrogenic gene expression in fetal chondrocytes transcriptomics data as well as single cell sequencing data when compared to adult chondrocytes. We also found CpGs losing methylation with age and genes associated with these CpGs showed upregulation in adult chondrocytes. miRNAs are known to play a key role in regulating chondrocyte development and homeostasis with age ^60^. In mammalian cells, DNA methylation is known to direct miRNA biogenesis ^61^. Hence, regulating expression of miRNAs by modulating DNA methylation may also act as a novel therapeutic strategy for chondrocyte repair and regeneration. We also observe gain of methylation in age-correlated CpGs for miRNAs known to be involved in chondrocyte homeostasis. Moreover, interrogation of the chromatin states for the age-correlated CpGs provided a clue towards enrichment of bivalent promoters during development. Enhancer chromatin states were also enriched across ontogeny providing a clue towards the region of transcriptional regulation.

STAT3, a key transcriptional factor, has been previously known to be involved in regulating stemness, development, and regeneration of tissues and organs. We have previously reported that STAT3 is highly expressed in anabolic fetal chondrocytes ^23^ and its involvement in chondrocyte development. Here, we observe that putative binding targets of STAT3 in fetal and adult chondrocytes are different and they are associated with distinct signaling pathways. We compared our results with transcriptomic data from fetal chondrocytes ^24^, STAT3 knocked down fetal chondrocytes ^49^ and chromatin accessibility data and found well known genes including ACAN, COL16A1, COL27A1, COL2A1, DUSP7, and KCNS1 to be the putative targets. Of these, age-correlated CpGs associated with COL2A1 was shown to gain methylation with age. In adult chondrocytes, of the 21 putative STAT3 targets, TLR1 and CD14 associated age-correlated CpGs were shown to lose methylation with age. STAT3 being a pleiotropic factor, regulates its targets in a context-specific manner. Thus, we also determined STAT3 targets in disease i.e., osteoarthritic chondrocytes and compared them to targets in development. In a nutshell, we observed the change in milieu of putative STAT3 targets in development and disease. Moreover, the critical role of STAT3 in development intrigued us to understand its effect in modulating DNA methylation. Genetic manipulation of STAT3 in fetal chondrocytes, induced a global hypermethylation, indicative of its role in maintaining an immature phenotype in chondrocytes.

The most challenging task in the field of aging is to determine a valid and reliable age predictor that will help understand how to slow, halt or even reverse aging ^62^. ‘Epigenetic clocks’ are accurate DNA methylation age estimators, which are built by regressing a transformed version of chronological age on a set of CpGs using a supervised machine learning model ^13^. In this study, we applied an epigenetic clock that is tailor-made for adult human chondrocytes and will be extremely useful in accurately estimating epigenetic age of adult chondrocytes. Our previous work has shown the importance of a small molecule STAT3 agonist that promotes cartilage repair and increases proliferation of chondrocytes ^23^. We used this chondrocyte clock, to explore the impact this small molecule has on epigenetic age in aged adult articular chondrocytes. Interestingly, we observed a decrease in epigenetic age in treated cells with a global hypomethylation in the genome.

In summary, the data presented here will serve as a foundation to understand the complex regulation of the epigenome across human chondrocyte ontogeny. Besides, it also provides strong evidence for the crucial role of STAT3 in modulating the epigenome during chondrocyte development. The novel epigenetic clock presented here will help researchers to capture pivotal aspects of biological age in adult chondrocytes. We anticipate this work will shed light towards chondrocyte aging with newer perspectives for development of rejuvenation agents.

## Methods

### Chondrocyte sample collection

Fetal tissue samples (14wks-19wks) were obtained from Novogenix Laboratories. All donated material was anonymous, carried no personal identifiers and was obtained after informed consent. Sex of the specimens was unknown. Adult human primary (21yrs-87yrs) and osteoarthritic tissue samples (55-60yrs) were obtained from National Disease Research Interchange (NDRI). Primary tissues were manually cut into small pieces and digested for 4–16 h at 37 °C with mild agitation in digestion media consisting of DMEM (Corning) with 10% FBS (Sigma), 1 mg/mL dispase (Gibco), 1 mg/mL type 2 collagenase (Worthington), 10 –µg/mL gentamycin (Teknova) and primocin (Invivogen).

### Cell culture and treatments

Only early passages of fetal and adult chondrocytes (P0) were used for experimentation to avoid de-differentiation and loss of cartilage phenotype^63^. Fetal and adult chondrocytes were cultured in DMEM F12 medium containing 10% (vol/vol) fetal bovine serum and 1% Penicillin-Streptomycin (vol/vol) at 37 °C in a humidified atmosphere of 95% air and 5% CO2. Media was replenished with DMEM F12 medium containing 1% (vol/vol) fetal bovine serum and 1% Penicillin-Streptomycin (vol/vol) once treatments were added.

Fetal chondrocytes were transduced with doxycycline inducible STAT3 shRNA or scrambled lentiviral particles (Dharmacon) and treated with Doxycycline every 48hrs. After 4 weeks of infection, transduced cells were sorted for RFP fluorescence.

Aged adult chondrocytes (55yrs-87yrs) were treated with or without a modified form of the small molecule STAT3 agonist, RCGD 423F N-(4-Fluorophenyl)-4-phenyl-2-thiazolamine; synthesized and provided by J-STAR Research at 10µM for 2weeks.

### FACS

FACS for fetal chondrocytes transduced with STAT3 shRNA or scrambled was performed on a BD FACSAria IIIu cell sorter. Cells were washed in 1% FBS and stained with DAPI for viability. Populations of interest based on DAPI negativity expression and RFP expression were directly sorted into DMEM/F12 containing 10% FBS with 1% P/S/A.

### RNA extraction and quantitative Real-Time PCR

Total RNA was extracted from live sorted fetal chondrocytes transduced with STAT3 shRNA or scrambled using the RNeasy Mini Kit (Qiagen). 500 ng of RNA was reverse transcribed using the Maxima First Strand cDNA Synthesis Kit (Thermo Fisher). Power SYBR Green (Applied Biosystems) RT-PCR amplification and detection was performed using an Applied Biosystems Step One Plus Real-Time PCR machine. The comparative Ct method for relative quantification (2-ΔΔCt) was used to quantitate gene expression, where results were normalized to Rpl7 (ribosomal protein L7). Primer sequences are available upon request. Results were analyzed using 2-tailed Student’s t test in GraphPad Prism 9.0.

### Genomic DNA extraction

Genomic DNA was extracted using QIAGEN DNeasy® Tissue kit or QIAamp® DNA Micro Kit depending on the starting number of cells. For DNeasy® Blood or Tissue kit samples were first lysed using Proteinase K. Lysate was loaded into the DNeasy Mini spin column and centrifuged to selectively bind DNA to the DNeasy membrane as contaminants pass through. Subsequent washing steps remove remaining contaminants and enzyme inhibitors. For QIAamp® DNA Micro Kit samples were lysed under high denaturing conditions at elevated temperatures in the presence of Proteinase K and Buffer ATL. Buffer AL was added to lysates followed by loading into QIAamp MinElute column and centrifugation. Residual contaminants or inhibitors are washed off using first Buffer AW1 and then Buffer AW2. Purified genomic DNA from either kit was eluted in water and quantified by Nanodrop confirming for high 260/280 purity ratio.

### DNA methylation data

The Illumina Infinium Methylation EPIC BeadChip array was used to perform DNA methylation profiling. This platform measures bisulfite conversion–based, single-CpG-resolution DNA methylation levels at 866,836 CpG sites in the human genome. Methylation levels are quantified by β values which is the ratio of intensities between methylated (signal A) and un-methylated (signal B) alleles. Specifically, the β value is calculated from the intensity of the methylated (M corresponding to signal A) and un-methylated (U corresponding to signal B) alleles, as the ratio of fluorescent signals β = Max(M,0)/[Max(M,0) + Max(U,0) + 100]. Thus, β values range from 0 (completely un-methylated) to 1 (completely methylated) ^64^.

### Analysis of Infinium EPIC methylation data

The R package “minfi” was used for analysis of the data ^65, 66^. Raw IDAT files were read and preprocessed and probes with high detection p-value (p-value>0.05) and potential SNP contamination were filtered. Normalization of data was done using the preprocessFunnorm function to generate Beta values per probe. Beta values provide the percentage of CpG methylation per probe with 0 being unmethylated and 1 fully methylated. Differentially methylated probes were identified by dmpFinder in logistic regression mode for appropriate contrasts followed by statistical analysis using an empirical Bayes method and then filtered by significance threshold (p-value<0.05, F-test). Annotation of probes was performed with the R package ‘IlluminaHumanMethylationEPICanno.ilm10b4.hg19’ version 0.6.0 for hg19 genome build. For EWAS approach ^55, 56, 67^, the DNA methylation changes were examined for association with chondrocyte age using the function “standardScreeningNumericTrait” from the “WGCNA” R package ^68^.

### DNA methylation age and Epigenetic clock

The chondrocyte clock was developed using both novel and existing methylation data from chondrocytes, cartilage and bone (Horvath 2021, in preparation). The age was regressed on DNA methylation levels using elastic net regression as implemented in the R function glmnet. The epigenetic clock for bones is described in separate article (Horvath 2021, in preparation).

### Bulk-RNA sequencing data analysis

Reads were aligned to human genome (hg19) using STAR aligner ^69^. Normalization was done using counts per million (CPM) method. Transcript levels were quantified to the reference using Partek E/M (build version 10.0.21.0210) with default parameters. Genes were considered to be differentially expressed based on fold change>1.5 and p-value<0.05. Gene set enrichment analysis was performed by Enrichr ^70–72^ using Fisher’s exact test (p-value<0.05). The background for enrichment was a lookup table of expected ranks and variances for each term in the library. These expected values were precomputed using Fisher’s exact test for many random input gene sets for each term in the gene set library.

### miRNA-sequencing and analysis

RNA was isolated using miRNeasy Micro Kit (Qiagen) according to manufacturer’s protocol. Briefly, samples were lysed by QIAzol lysis reagent followed by addition of chloroform and centrifugation to separate the solution into phases. The upper aqueous phase was extracted, ethanol was added, and samples were loaded into RNeasy MinElute spin column. Thereafter a specialized protocol was used to separate the enriched miRNA fraction. miRNA was quantified using Qubit fluorometer, and run on Agilent Bioanalyzer 2100 for quality control. Libraries were prepared using NEBNext Multiplex Small RNA Library Prep Set (Illumina) according to the manufacturer’s protocol. The workflow consists of adapter ligation, cDNA synthesis, PCR enrichment, clean up and size selection. Different adapters were used for multiplexing samples in one lane. Sequencing was performed on Illumina HiSeq 2500 with single-end 50 base pair reads. Reads were aligned to human genome (hg38) using Bowtie ^73^. Normalization was done using counts per million (CPM) method and miRNAs levels were quantified (miRBasev22). A lognormal with shrinkage model was used for differential expression analysis. miRNAs were considered to be differentially expressed based on fold change>1.5 and p-value<0.05.

### Single-cell sequencing using 10X Genomics

Single cell samples were prepared using Single Cell 3^/^ Library & Gel Bead Kit v2 and Chip Kit (10X Genomics) according to the manufacturer’s protocol. Briefly samples were FACS sorted using DAPI to select live cells followed by resuspension in 0.04% BSA-PBS. Nearly 1,200 cells/µl were added to each well of the chip with a target cell recovery estimate of 8,000 cells. Thereafter Gel bead-in Emulsions (GEMs) were generated using GemCode Single-Cell Instrument. GEMs were reverse transcribed, droplets were broken and single stranded cDNA was isolated. cDNAs were cleaned up with DynaBeads and amplified. Finally, cDNAs were ligated with adapters, post-ligation products were amplified, cleaned up with SPRIselect. Purified libraries were submitted to UCLA Technology Center for Genomics & Bioinformatics for quality check and sequencing. The quality and concentration of the purified libraries were evaluated by High Sensitivity D5000 DNA chip (Agilent) and sequencing was performed on NextSeq500.

### 10X sequencing data analysis

Raw sequencing reads were processed using Partek Flow Analysis Software (build version 10.0.21.0210). Briefly, raw reads were checked for their quality and trimmed. Reads with an average base quality score per position >30 were considered for alignment. Trimmed reads were aligned to the human genome version hg38-Gencode Genes-release 30 using STAR -2.6.1d with default parameters. Reads with alignment percentage >75% were de-duplicated based on their unique molecular identifiers (UMIs). Reads mapping to the same chromosomal location with duplicate UMIs were removed. Thereafter ‘Knee’ plot was constructed using the cumulative fraction of reads/UMIs for all barcodes. Barcodes below the cut-off defined by the location of the knee were assigned as true cell barcodes and quantified. Further noise filtration was done by removing cells having >3% mitochondrial counts and total read counts >24,000. Genes not expressed in any cell were also removed as an additional clean-up step. Cleaned up reads were normalized using counts per million (CPM) method followed by log transformation generating count matrices for each sample. Samples were batch corrected on the basis of expressed genes and mitochondrial reads percent. Dotplot was generated in R (v4.0.3) using ggplot2 (v3.3.3) package.

### ChromHMM analysis

We conducted ChromHMM ^74^ chromatin state enrichment analysis with chromatin state annotations from Fetal 17 weeks and Adult chondrocytes tissues using a previously defined 12- state model ^24^. Hypermethylated and hypomethylated age-correlated CpGs were determined by EWAS as mentioned previously. Using the OverlapEnrichment command of ChromHMM we computed the enrichment for the coordinates set of hypermethylated and hypomethylated age-correlated CpGs. We did the same for the coordinates of all CpGs on the array, and then divided the hypermethylated and hypomethylated age-correlated CpG enrichment values by these enrichment values to obtain the enrichment relative to the array background.

### ATAC-sequencing and data analysis

Samples were washed, lysed followed by nuclei tagmentation and adapter ligation by Tn5 using the Nextera DNA Sample Preparation kit (Illumina). Transposed DNA fragments were amplified using the NEBNext Q5 HotStart HiFi PCR Master Mix with regular forward and reverse barcoded primers. The final product was purified with MinElute PCR Purification kit (Qiagen), and quality checked on 2100 Bioanalyzer (Agilent). Sequencing was performed on Illumina HiSeq 2500 with single-end 50 base pair reads. The initial quality of the raw fastq files were checked using FastQC (https://www.bioinformatics.babraham.ac.uk/projects/fastqc/). Reads were trimmed using Cutadapt v2.10 ^75^ in paired-end mode. Trimmed reads were aligned to human genome build hg19 using bowtie2. PCR duplicates were removed from the aligned reads followed by sorting and indexing of the bam files by SAMtools v1.11 ^76^. Bam coverage maps were generated using bamCoverage from the deepTools suite v3.5.0 ^77^. Significant peaks (p-value<0.05) were called using MACS2 ^78^ and annotated using R.

### Cleavage Under Targets and Release Using Nuclease (CUT&RUN)

In situ chromatin profiling using CUT & RUN was performed according to Skene et al ^34^. Briefly samples were FACS sorted using DAPI to select live cells and 10,000 cells were collected in 10%FBS-PBS media. Cell nuclei were immobilized on Concanavalin A beads after washing. pSTAT3 (Tyr705,D3A7,9145,Cell signaling technology) or normal rabbit IgG antibodies (3900,Cell signaling technology) were incubated with the nuclei overnight in the presence of 0.02% digitonin at 4°. The next day, 700ng/mL of proteinA-micrococcal nuclease (pA-Mnase purified in house with vector from Addgene 86973 ^79^) were incubated with the nuclei at 4 degrees for an hour. After washing, the tubes were placed in heat blocks on ice set to 0 degrees, CaCl_2_ (1mM) was added and incubated for 30 min before 2X Stop buffer containing EDTA was added. DNA was extracted using Qiagen DNA isolation kit according to manufacturer’s protocol Purified DNA was quantified in Qubit and Bioanalyzer (2100) traces using D5000 high sensitivity chip were run to determine the size of the cleaved products. UMI-coded libraries were generated using Swift Biosciences-ACCEL-NGS® 2S PLUS DNA LIBRARY KITS according to manufacturer’s protocol. Pair-end (75bp) Illumina sequencing was performed on the UMI-coded and amplified libraries using NextSeq platform.

### CUT&RUN data analysis

UMI-tools ^80^ ‘extract’ function was used to remove UMIs from each read of the raw fastq files and append them to the read name. The initial quality of the raw fastq files were checked using FastQC (https://www.bioinformatics.babraham.ac.uk/projects/fastqc/). Reads were trimmed using Cutadapt v2.10 ^75^ in paired-end mode. Trimmed reads were aligned to human genome build hg19 using bowtie2. Next, aligned reads were deduplicated and PCR duplicates were removed by UMI-tools ‘dedup’ function followed by sorting and indexing of the deduplicated bam files by SAMtools v1.11 ^76^. Bam coverage maps were generated using bamCoverage from the deepTools suite v3.5.0 ^77^. Heatmaps were generated using computeMatrix and plotHeatmap from the deepTools suite v3.5.0. Significant peaks (p-value<0.05) were called from the deduplicated reads using MACS2 ^78^ and annotated using the R package ChIPseeker ^81^. Subsequently, peak files were used to determine enriched motifs using HOMER v4.11.1 ^82^. Functional enrichment analysis for the nearest genes annotated to the peaks was determined by the R package clusterProfiler ^83^. Two-way Venn diagrams were generated using BioVenn ^84^, while 4-way Venn diagrams were constructed using InteractiVenn ^85^.

### Western blot analysis

Osteoarthritic chondrocytes were lysed in RIPA Lysis and Extraction Buffer (Pierce) containing protease inhibitors (Pierce) followed by sonication with a 15-second pulse at a power output of 2 using the VirSonic 100 (SP Industries Company). Protein concentrations were determined by BCA protein assay (Pierce) and boiled for 5 minutes with Laemmli Sample Buffer (Bio-Rad, Hercules, CA). Proteins were separated on acrylamide gels and analyzed by Western blot using primary antibodies: anti-pSTAT3 (9145) and anti-Histone H3 (9515; all from Cell Signaling). Histone H3 antibody was used as a loading control. Proteins were resolved with SDS-PAGE utilizing 4–15% Mini-PROTEAN TGX Precast Gels and transferred to Trans-Blot Turbo Transfer Packs with a 0.2-µm pore-size nitrocellulose membrane. The SDS-PAGE running buffer, 4–15% Mini-PROTEAN TGX Precast Gels, Trans-Blot Turbo Transfer Packs with a 0.2-µm pore-size nitrocellulose membrane were purchased from Bio-Rad. Nitrocellulose membranes were blocked in 5% nonfat milk in 0.05% (v/v) Tween 20 (Corning). Membranes were then incubated with primary antibodies overnight. After washing in PBS containing 0.05% (v/v) Tween 20 (PBST), membranes were incubated with secondary antibodies (31460 and 31430, Thermo Scientific). After washing, development was performed with the Clarity Western ECL Blotting Substrate (Bio-Rad).

### Data Availability

All data is deposited in GEO and is available under the accession number GSEXXXX.

---

**Figure S1.**
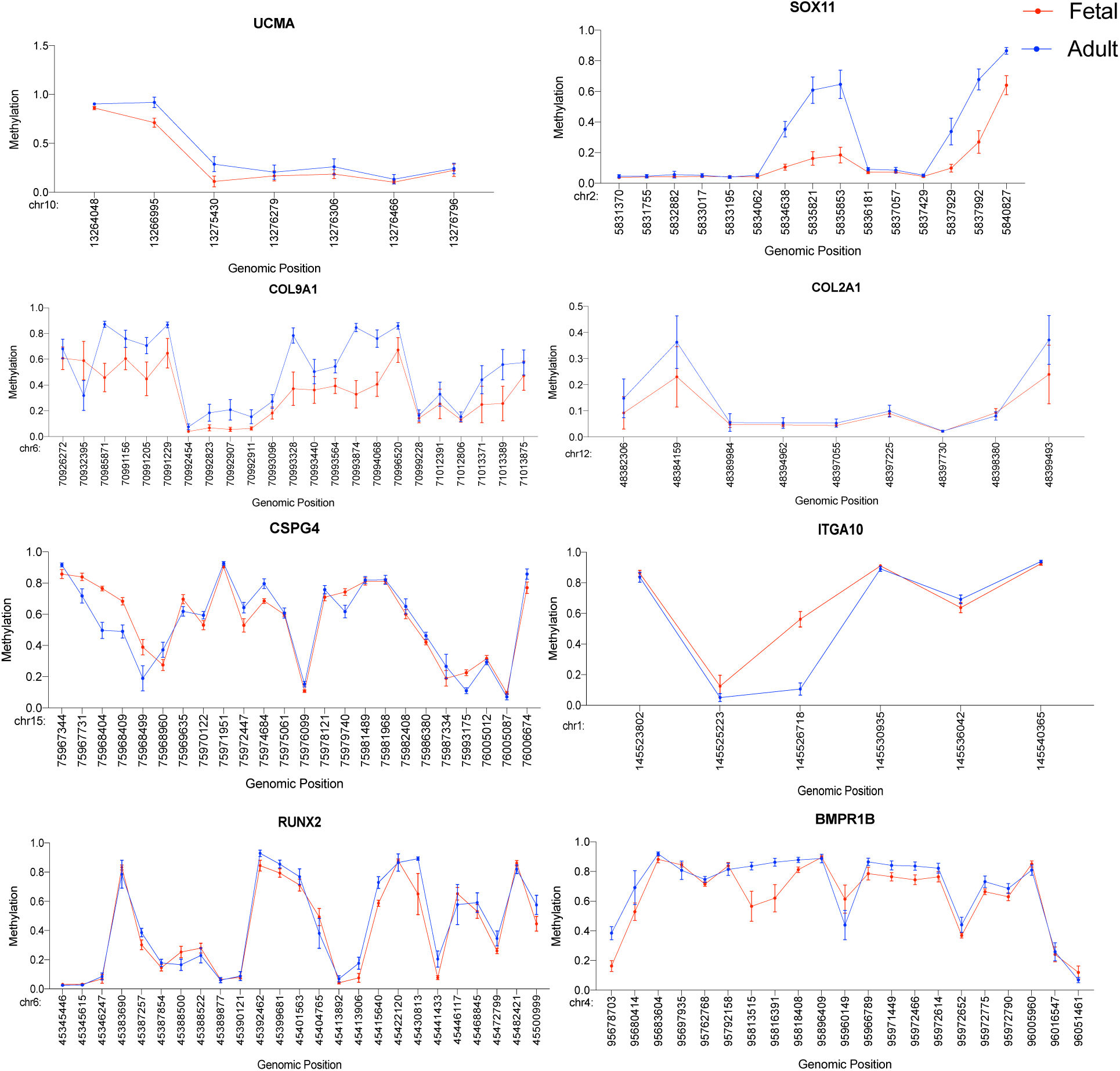
Gain of methylation in age-correlated CpGs associated with chondrogenic genes. Scatterplot showing the methylation level and genomic coordinates for all age-correlated CpGs associated with the chondrogenic genes. Mean with standard deviation is plotted.

**Figure S2.**
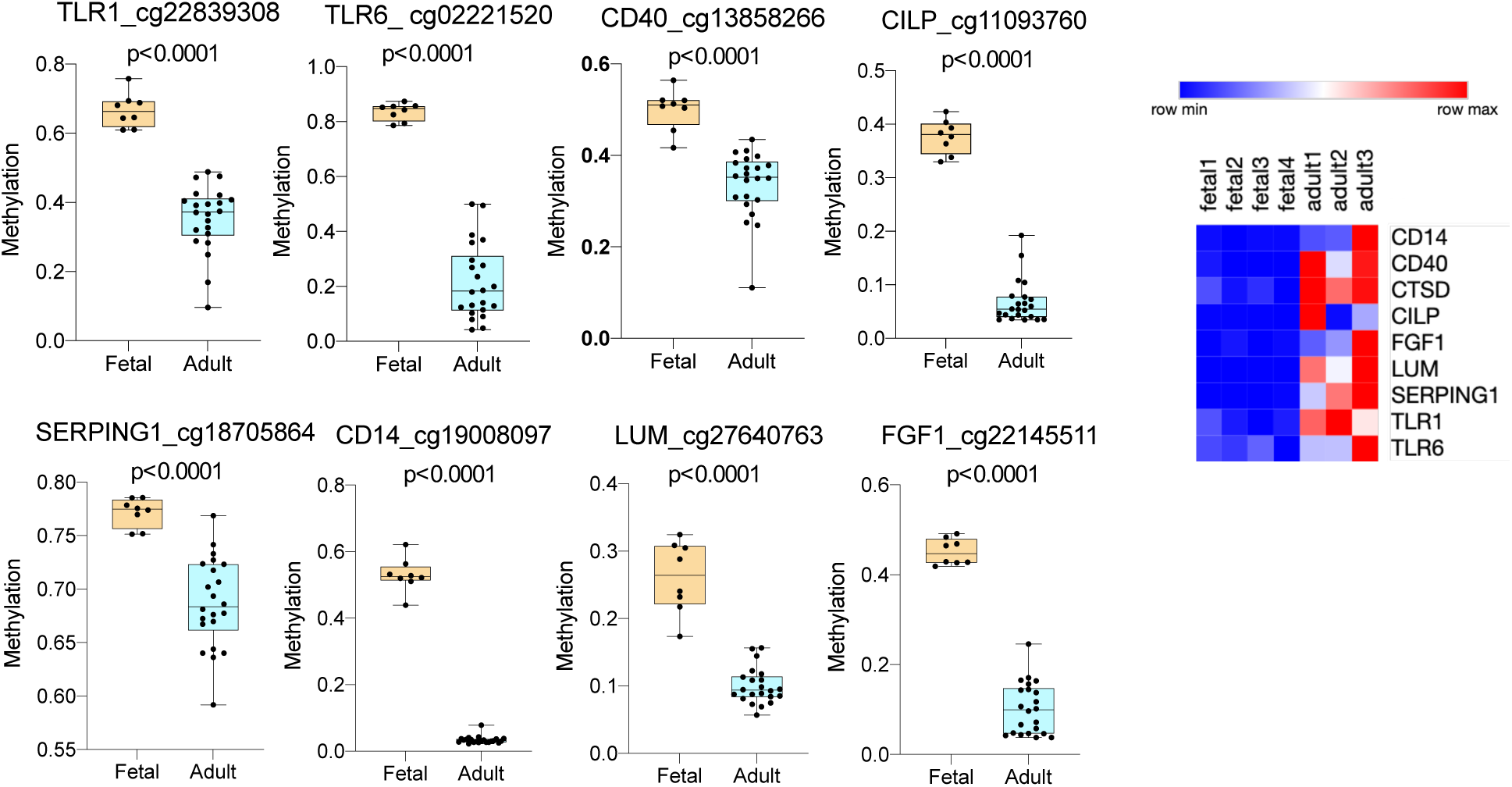
Loss of methylation in age-correlated CpGs. Boxplot showing methylation level of representative age-correlated CpGs (i.e., CpGs with highest hypomethylation change across age). Transcriptomic profile for the genes in fetal and adult chondrocytes is also shown. Hinges of all boxplots extend from the 25th to 75th percentiles. The line in the middle of the box is plotted at the median. P-values are calculated using 2-tailed Student’s t test.

**Figure S3.**
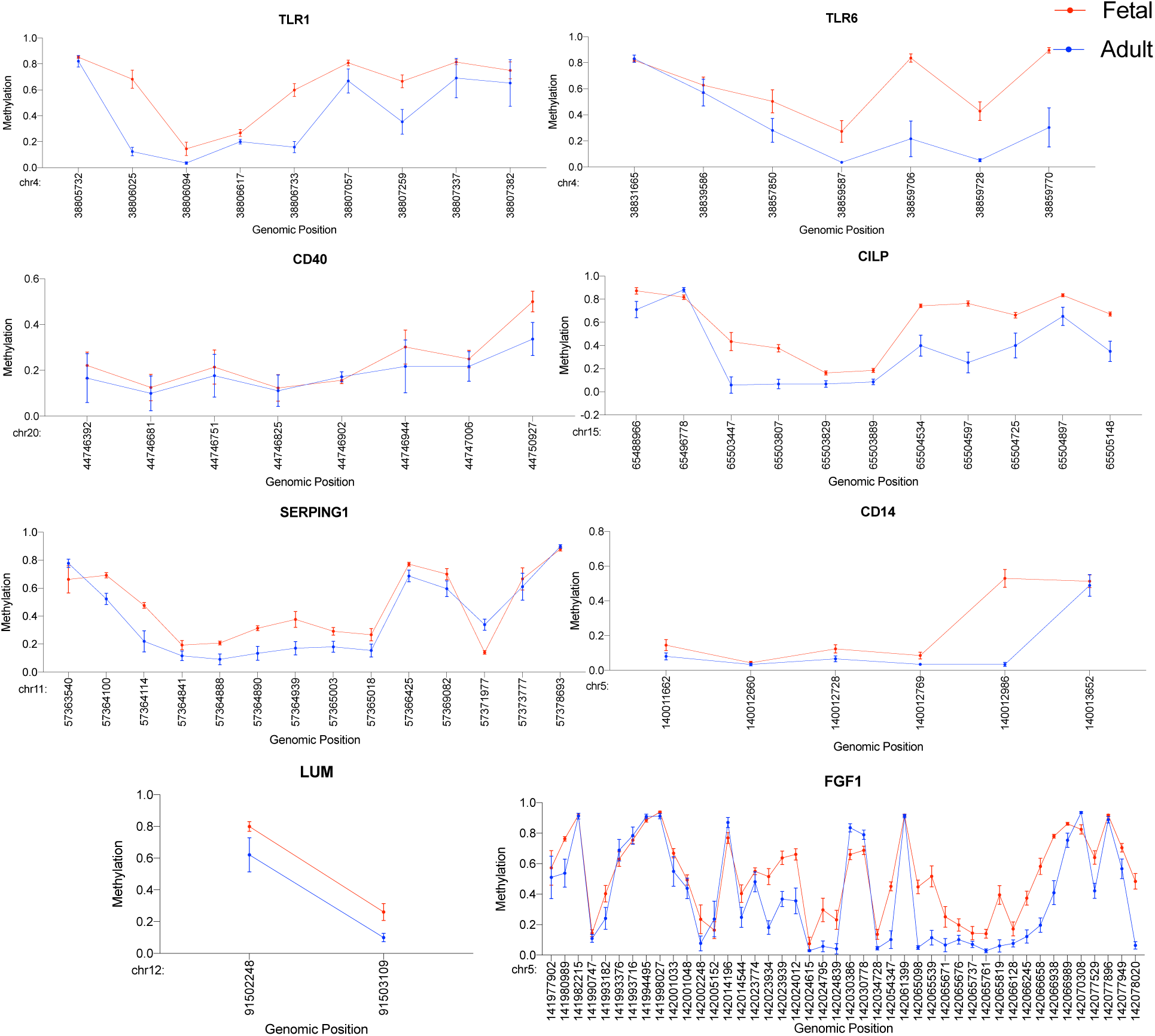
Scatterplot showing methylation level and genomic coordinates for all age-correlated CpGs losing methylation with age. Mean with standard deviation is plotted.

**Figure S4.**
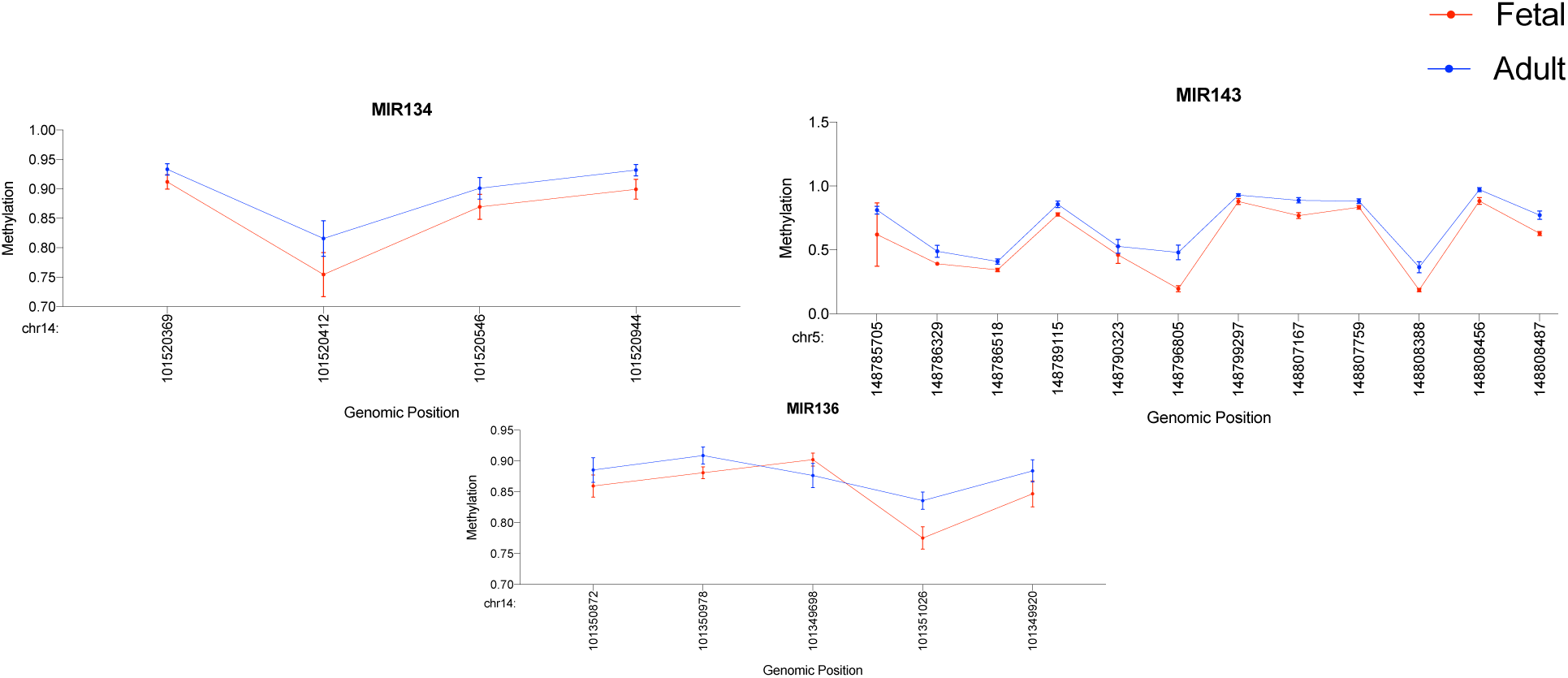
Gain of methylation in age-correlated CpGs associated with miRNAs. Scatterplot showing the methylation level and genomic coordinates for all age-correlated CpGs associated with the miRNAs. Mean with standard deviation is plotted.

**Figure S5.**
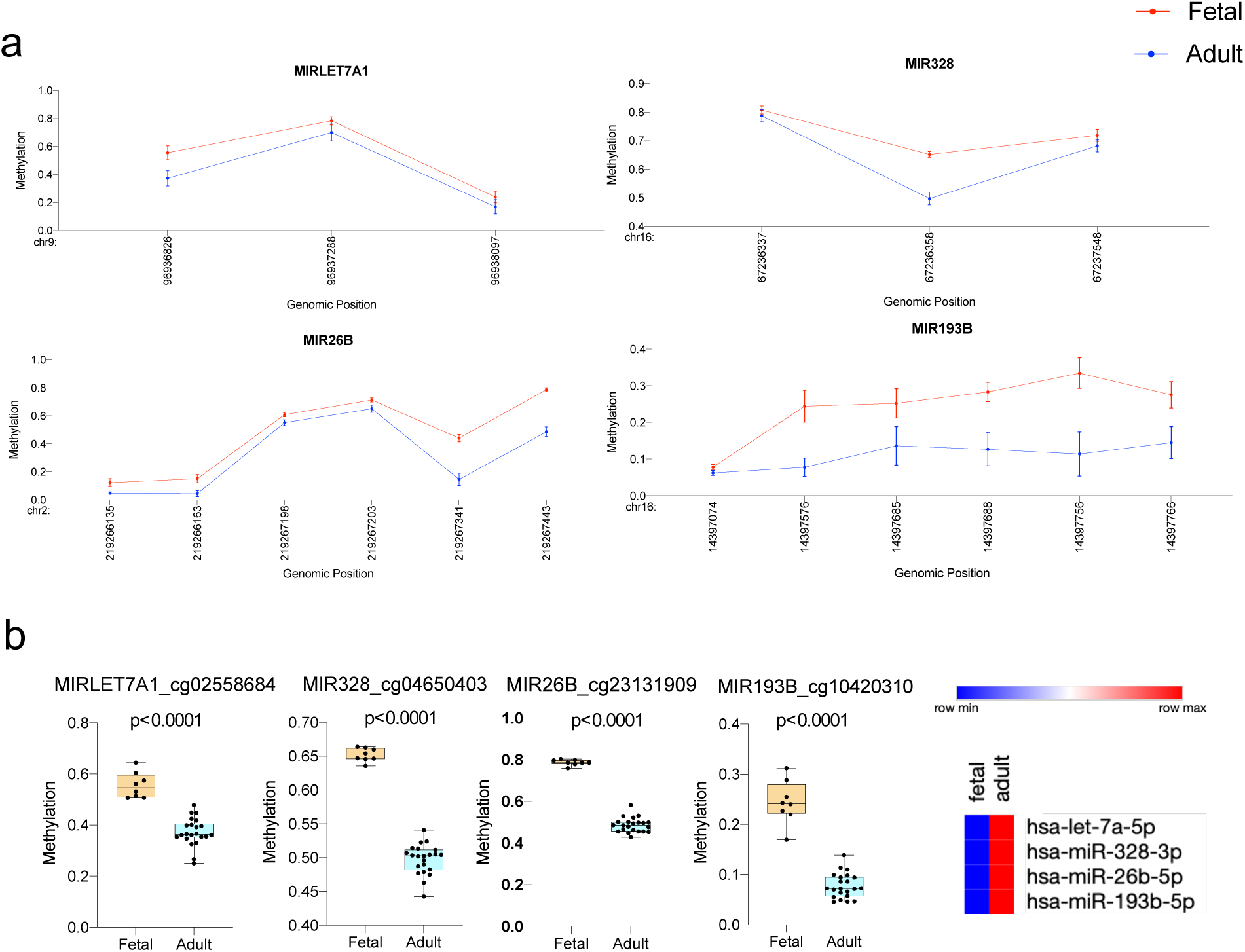
Loss of methylation in age-correlated CpGs associated with miRNAs. **a.** Scatterplot showing the methylation level and genomic coordinates for all age-correlated CpGs associated with the miRNAs. Mean with standard deviation is plotted. **b.** Boxplot showing methylation level of representative age-correlated CpGs (i.e., CpGs with highest hypomethylation change across age). miRNA expression profile in fetal and adult chondrocytes is also shown. Hinges of all boxplots extend from the 25th to 75th percentiles. The line in the middle of the box is plotted at the median. P-values are calculated using 2-tailed Student’s t test.

**Figure S6.**
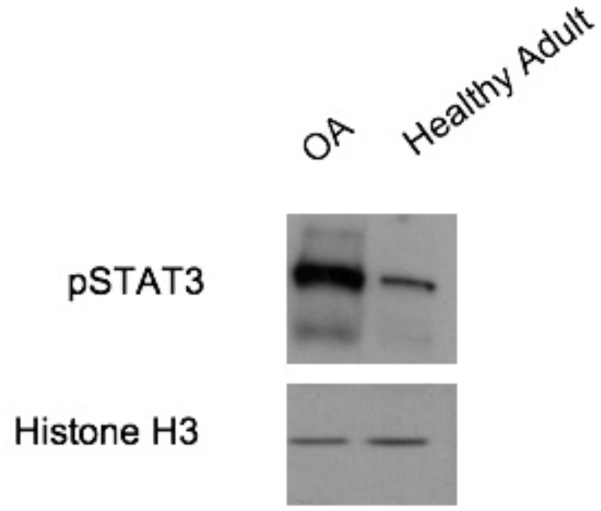
Active STAT3 is highly expressed in osteoarthritic chondrocytes as compared to healthy adult chondrocytes.

**Figure S7.**
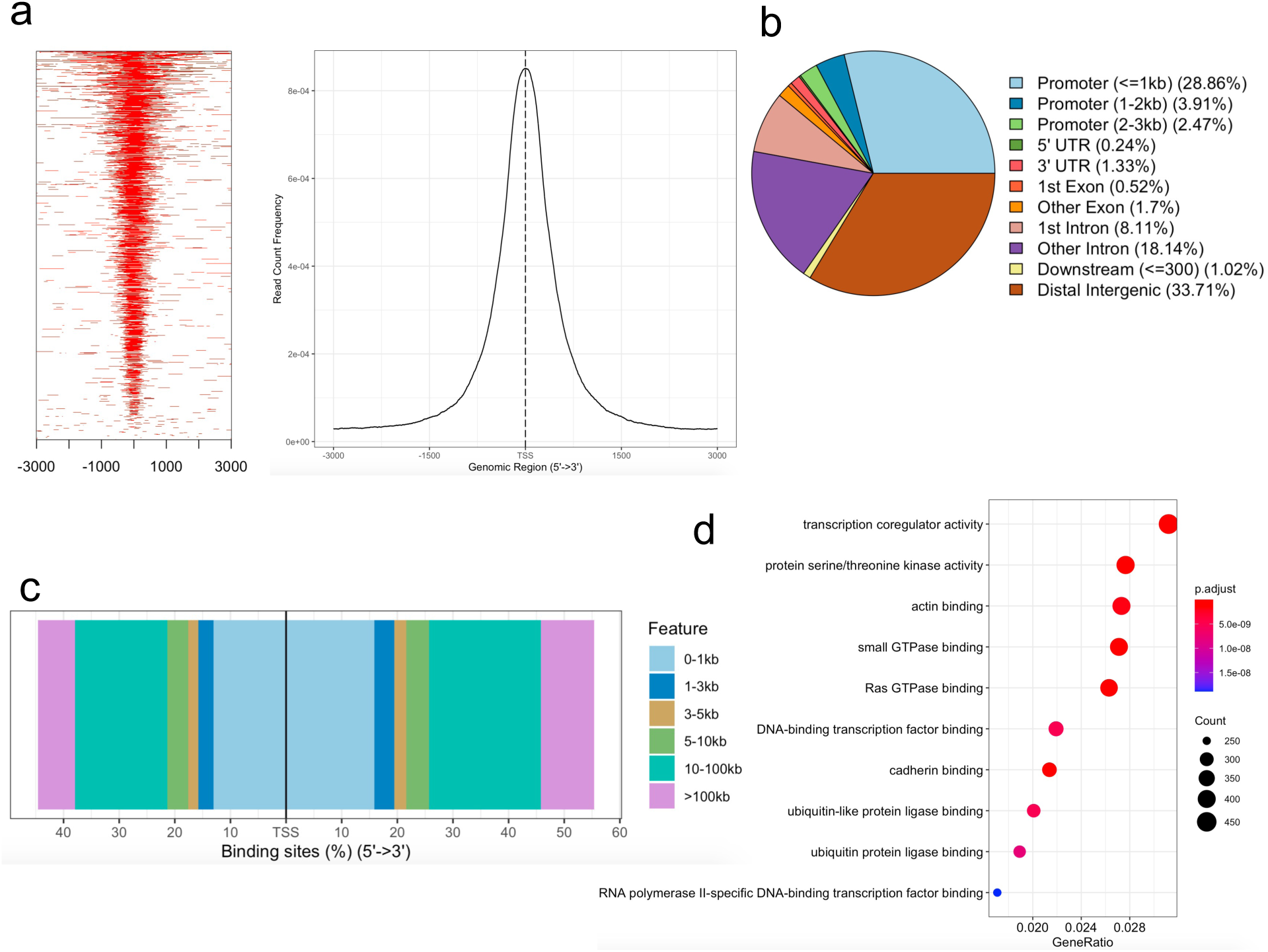
ATAC-Seq for fetal chondrocyte **a.** Heatmap showing enrichment of reads and distribution of peaks across ±3kb of TSS. **b,c.** Pie chart and bar plot showing the distribution of genomic features. **d.** Functional enrichment of target genes. P-values were adjusted using Benjamini-Hochberg correction method.

**Figure S8.**
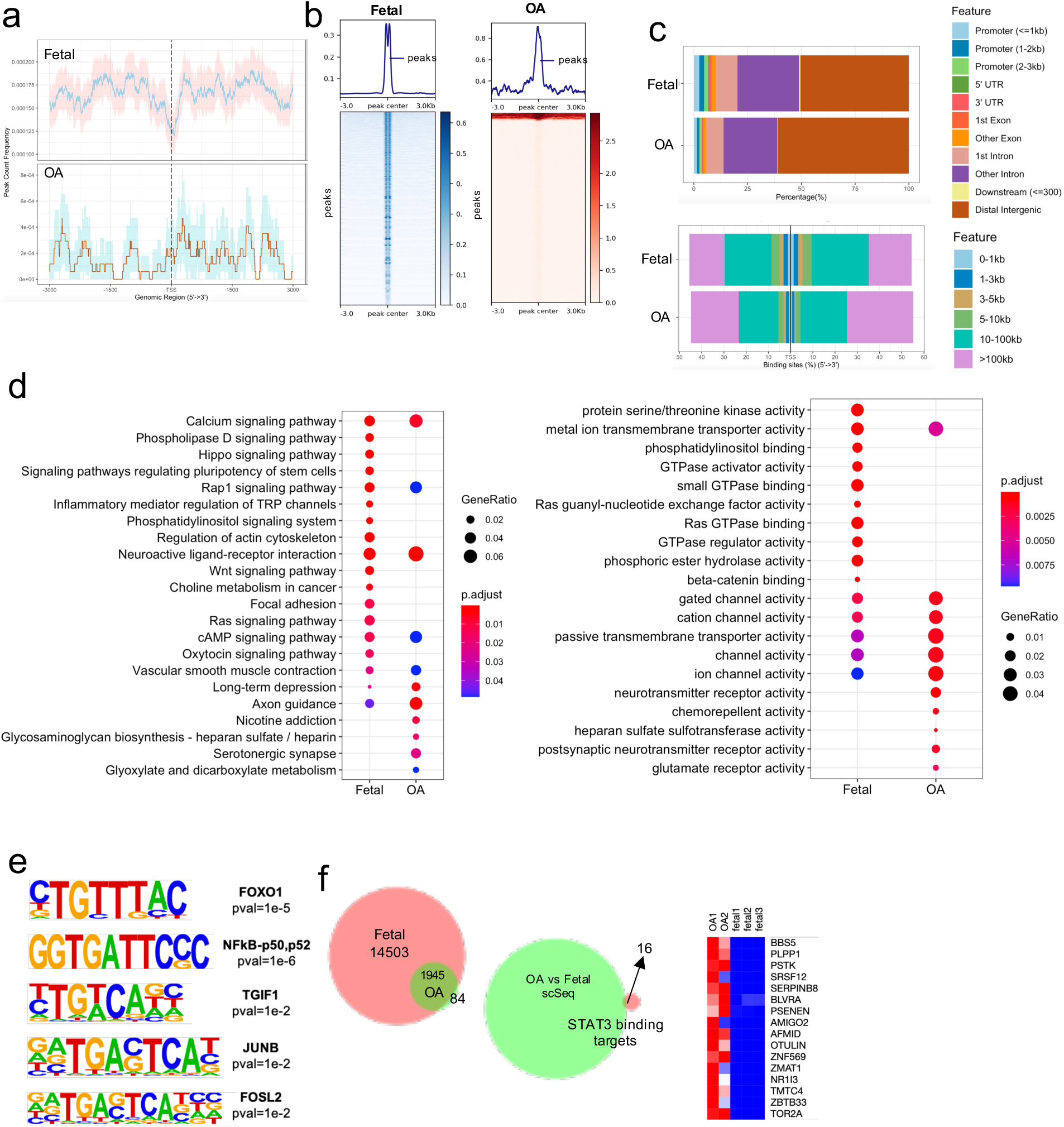
STAT3 binding targets in disease. OA=osteoarthritis **a.** Distribution of peak count frequency across ±3kb of TSS. Confidence intervals shown by the shadows following each curve were estimated by bootstrap method using 500 iterations **b.** Heatmap showing enrichment of reads in peak summits. **c.** Bar plot showing the distribution of genomic features for peaks in fetal and osteoarthritic chondrocytes. **d.** Gene enrichment analysis of putative STAT3 target genes. P-values were adjusted using Benjamini-Hochberg correction method. **e.** DNA motif enrichment analysis for putative STAT3 binding targets. Binomial distribution was used to score motifs. **f.** Venn diagram showing the overlap between putative STAT3 targets in fetal and osteoarthritic chondrocytes. 84 exclusive fetal chondrocyte targets were overlapped with OA vs fetal single cell sequencing data. Heatmap shows the expression profile of the 16 final targets obtained for osteoarthritic chondrocytes.

**Figure S9.**
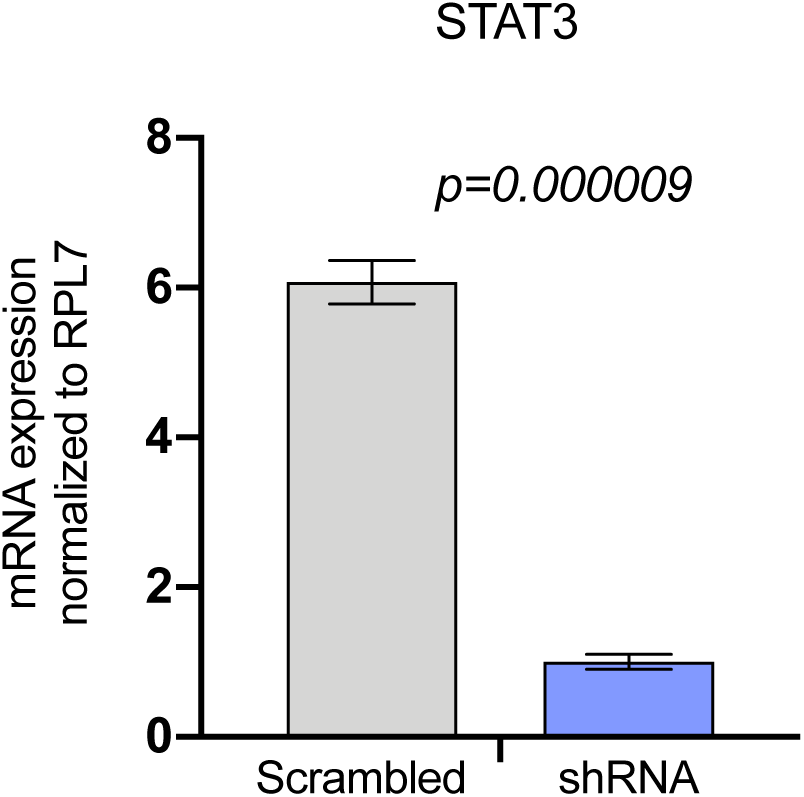
qRT-PCR data analysis for STAT3 in scrambled and STAT3 shRNA fetal chondrocytes. Statistical analysis was performed using 2-tailed Student’s t test in GraphPad Prism 9.0 and p-value <0.05 was considered as statistically significant. Mean with standard deviation is plotted.

